# Protein corona modulates interaction of spiky nanoparticles with lipid bilayers

**DOI:** 10.1101/2021.05.27.446016

**Authors:** Jean-Baptiste Fleury, Marco Werner, Xavier Le Guével, Vladimir A. Baulin

**Affiliations:** Experimental Physics and Center for Biophysics, Universitat des Saarlandes, 66123 Saarbruecken, Germany; Leibniz-Institut für Polymerforschung Dresden e.V., 16 Hohe Strasse 6, 01069 Dresden, Germany; Cancer Targets & Experimental Therapeutics, Institute for Advanced Biosciences (IAB), University of Grenoble Alpes - INSERM U1209 - CNRS UMR 5309- 38000 Grenoble, France; Departament Química Física i Inorgànica, Universitat Rovira i Virgili, Marcel.lí Domingo s/n, 43007 Tarragona, Spain

## Abstract

The impact of protein corona on the interactions of nanoparticles (NPs) with cells remains an open question. This question is particularly relevant to NPs which sizes, ranging from tens to hundreds nanometers, are comparable to the sizes of most abundant proteins in plasma. Protein sizes match with typical thickness of various coatings and ligands layers, usually present at the surfaces of larger NPs. Such size match may affect the properties and the designed function of NPs. We offer a direct demonstration of how protein corona can dramatically change the interaction mode between NPs and lipid bilayers. To this end, we choose the most extreme case of NP surface modification: nanostructures in the form of rigid spikes of 10-20 nm length at the surface of gold nanoparticles. In the absence of proteins we observe the formation of reversible pores when spiky NPs absorb on lipid bilayers. In contrast, the presence of bovine serum albumin (BSA) proteins adsorbed at the surface of spiked NPs, effectively reduce the length of spikes exposed to the interaction with lipid bilayers. Thus, protein corona changes qualitatively the dynamics of pore formation, which is completely suppressed at high protein concentrations. These results suggest that protein corona can not only be critical for interaction of NPs with membranes, it may change their mode of interaction, thus offsetting the role of surface chemistry and ligands.

## Introduction

Gold nanoparticles (NPs) are used in many biomedical applications ranging from drug delivery^1,2^ and medical imaging^3^ to cancer treatment.^4–6^ The size, shape and surface of gold NPs define most of their physico-chemical properties.^7,8^ The behavior of gold NPs is well controlled by their physico-chemical properties. While NPs are dispersed in biological fluids, their behavior is more ambiguous.^9^ Proteins and biopolymers, abundantly present in plasma and other biological fluids, tend to aggregate and adsorb at NPs surface, forming a layer of water-soluble proteins (or biopolymers), called protein corona (PC).^10,11^ This protein layer is not static, but dynamically exchanging proteins with the bulk while its complex kinetics depends on the size and the structure of proteins present in the bulk.^12^ As a result, PC may change properties of NPs, for example, PC can suppresses most of the functional properties of ligand-functionalized well-controlled NPs and even trigger immune reaction.^13^ Thus, PC may screen, alter and suppress the contribute prepared in the lab and validated *in vitro*, NPs may not work *in vivo*: more than 95 % of the administered NPs end up at sites other than targeted tumors.^14^ Thus, PC plays an important role in cellular uptake, targeting, clearance and possible nanotoxicity of these NPs.^15,16^

Many strategies have been adopted to prevent the formation of PC and their interactions with the bio/nano-interface during the last decade.^17^ The difficulties reside in the dynamic nature of PC and that in biological media it consists of a myriad of different biomolecules.^15–17^ Large number of proteins, lipids or ligands (such as carbohydrates) that can cover NPs to prevent the PC formation were explored. ^10,18,19^ However, prevention of PC may also alter or inhibit the NPs functionality. ^10,18,19^ Some of these strategies even lead to paradoxical results, for example, coating NPs surfaces with hydrophilic polymers can prevent proteins from adsorbing, but, in turn, it can enhance the recognition of the NPs by the immune system. ^17,20,21^

It is usually considered that the effect of PC is directly proportional to the NP surface area available for the adsorption, ^22,23^ while the surface area strongly depends on the size and shape of NPs. With this, apart from the surface modification strategy, the shape of NPs has been also considered as a possible way to modulate the PC formation. Recent results show that rod-like NPs can accommodate significantly larger amounts of serum proteins compared to spherical NPs or specific nanostar-shaped NPs. ^22,23^

To demonstrate significant change of properties of NPs due to adsorption of proteins and even qualitative change of the behavior when interacting with lipid bilayers, in this manuscript we adopt the most radical form of surface modification of NPs, namely NPs with the shape in form of rigid spikes, so called spiky nanoparticles (SNs). These objects are of particular importance for interaction with proteins, since the sizes of spikes of few nanometers are of the order of magnitude of the sizes of most proteins in plasma.^24,25^ Thus, these objects are at the crossover between large NPs covered with protein corona and small NPs or nanoclusters with high curvatures and sizes comparable to protein sizes. Due to a particular shape, SNs have recently attracted interest as potential candidate to prevent the PC formation^23,26^ and the shape of spikes can be very broad. ^23,26,27^ This makes SNs the ideal objects to study the effects of adsorbed proteins also in other systems. It is noteworthy, that SNs have the shape similar viral particles.^28,29^ Surprisingly, viruses share many biophysical properties with artificial SNs in extracellular environments. ^29^ In particular, PC formation is occurring on the viral particle surface and this layer is critical for viral-host interactions. ^29^ Thus, it is interesting to investigate how the shape of a SN could prevent, or affect, the formation of a protein corona on the viral particle surface as well as study interaction of NPs with model cell membranes.^30,31^

In the first section we describe the physico-chemical properties and the geometry of SNs. Then we study the interaction of SNs of different shapes and sizes with lipid bilayers. The spikes at the surfaces of SNs which are comparable in length with the thickness of the bilayers induce pores in the bilayers, which are studied with patch-clamp technique and theoretically with bond-fluctuation model within Monte Carlo simulations. Next chapter describes the effect of protein corona adsorbed on SNs, which is manifested in the effective screening of spikes length, that change qualitatively the behavior of SNs and suppresses pore formation.

## Characterization of spiky nanoparticles

SNs were prepared by a standard seed-mediated growth method ^32^ starting from either 3 nm (Types A and B) or 15 nm (Types C and D) gold seeds (see Materials and Methods for details). This protocol results in SNs with gradually changing sizes and variable spike numbers from Type A to D. We chose this synthesis protocol because it provides SNs at a high yield, high monodispersity and with a precise control of spike numbers and spike lengths.^33^ The sizes and geometry of the spikes were determined from TEM images at the same magnifications for a large number of SNs (see Fig. 1A-D and additionally Fig. S1-S4). All dimensions of the four types of SNs are reported in Table 1. Full size corresponding to the average size of SNs including the metal core and the spikes.

**Figure 1:**
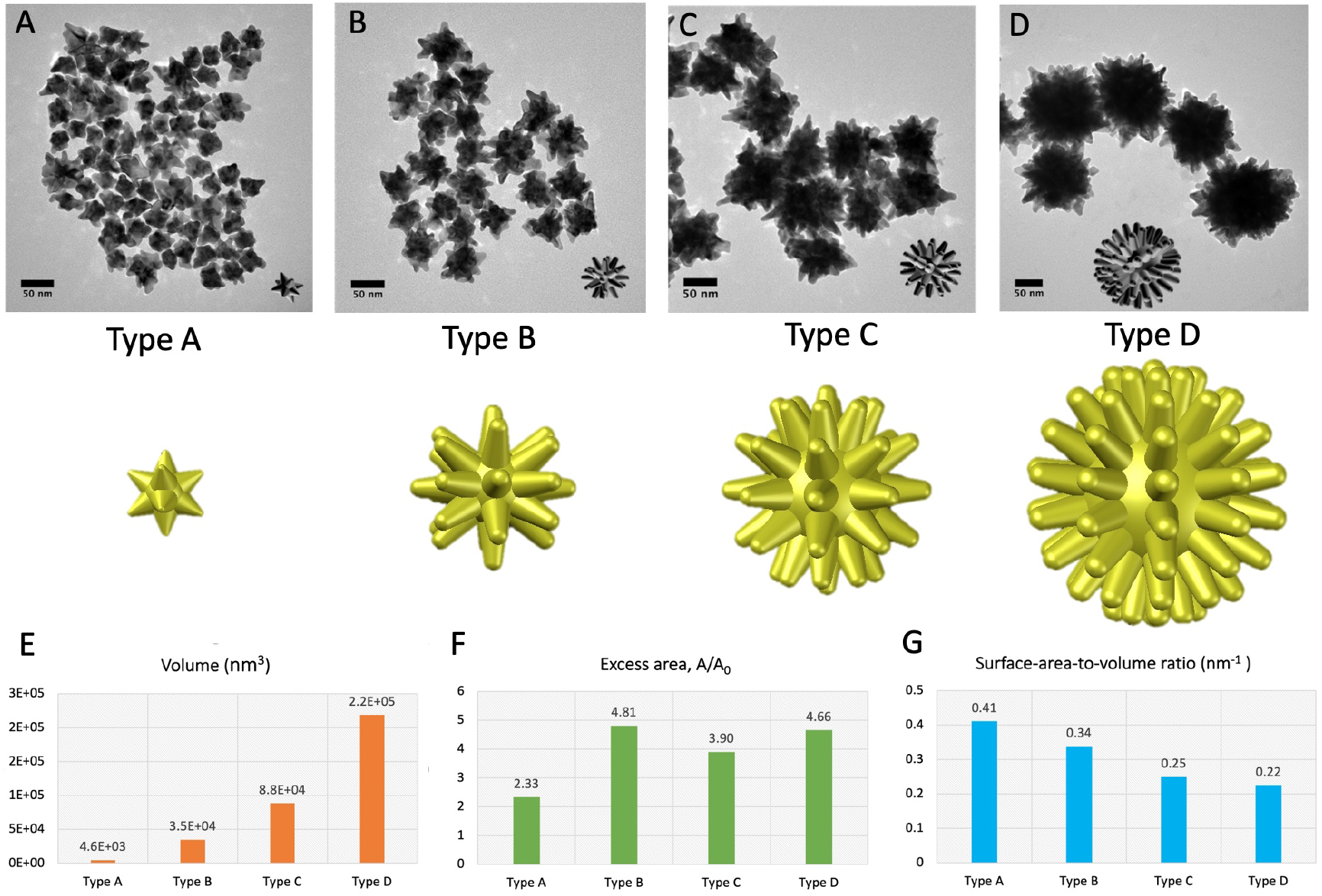
A), B), C), D): TEM images of SNs of types A, B, C, D and their 3D reconstructions, correspondingly. E) The calculated volume of each 3D model of SNs, F) Excess area of each 3D model, the total surface area, divided by the projected area, *A/A*_0_. G) Surface-area-to-volume ratio of SNs of types A, B, C, D.

**Table 1:** Measured geometrical parameters of SNs obtained from an average of 100 NPs TEM images.

|  | Core size (nm) | Full size (nm) | Number of spikes | Spike length (nm) | Spike angles (°) | Radius for spikes angle (°) | Angles between spikes (°) | Distance between spikes (nm) | Spikes thickness at half-height (nm) |
| --- | --- | --- | --- | --- | --- | --- | --- | --- | --- |
| Type A | 16± 2 | 44±4 | 7±2 | 9.7±2 | 51.6±1 | 5.1±1 | 120.8±18 | 23.4±5 | 9±1 |
| Type B | 27.8±4 | 66.5±9 | 10±2 | 17.4±4 | 35.4±7 | 5.4±2 | 71.9±15 | 21.8±4 | 10.2±1 |
| Type C | 42.4±4 | 103.9±12 | 14±2 | 18.2±3 | 40.1±9 | 6.2±2 | 87.6±20 | 19±3 | 11.1±2 |
| Type D | 58±5 | 140.9±8 | 19±3 | 21±4 | 38.7±7 | 5.9±2 | 60.5±15 | 17.4±3 | 12.1±1 |

SNs of Type A and B were obtained from 3 nm seeds, resulting in full sizes of 44 ± 4.7 nm for Type A and 66.5 ± 8.9 nm for Type B (Fig. 1 and Table 1). SNs of Type A are not only smaller, they have less spikes in average 7 ± 2 compared to 10 ± 2 for Type B and smaller lengths of the spikes, 9.7 ± 2.0 nm compared to 17.4 ± 4.0 nm for Type B. The angle between spikes is smaller (spikes are sharper) for Type A, 51.6 ± 1.0 compared to 35.4 ± 6.9 for Type B.

SNs of Type C and Type D were obtained from the 15 nm gold seeds and result in full sizes of 103 ± 11.9 nm and 140.9 ± 7.7 nm, respectively. We observed an increase of spike numbers but no significant changes of spike lengths from Type C to D. The angle between spikes and the distance between spikes have similar values as SNs of Type B. Noteworthy, that gold core has constantly increased for all types of SNs during the synthesis. This could be explained by the reaction of gold salt that reacted on gold seeds to produce bigger sizes and also to form spikes due to the presence of PVP.^34^ In addition, we observed that SNs exhibit a broad near-infrared plasmon band typical of anisotropic gold NPs^32^ (Fig. S5). The synthesized SNs were functionalized with thioctic bidentate sulfobetaine zwitterionic molecule (see Materials and Methods) to prevent aggregation and insure colloidal stability in the solution. Non-homogeneous distribution of the coating results in hydrophilic interiors of the SN between the spikes and hydrophobic tips of the spikes. The hydrophobicity of the tips allows for spikes to anchor the lipid bilayer and affects the protein adhesion locally.

The detailed geometrical parameters of each type of SNs obtained from TEM images, allow for the 3D reconstruction of an average SN of each type, as illustrated in Fig. 1A-D. The reconstruction, in turn, allows to estimate additional geometrical parameters: the total volume of an average SN of each type is gradually growing from Type A to D, Fig. 1E; the excess area, defined as the total area of SN divided by the projected area of the core, is smaller for Type A, while for B, C, D has similar values, Fig. 1F; the surface-area-to-volume ratio is gradually decreasing from Type A to D, Fig. 1G. This data is further used for Monte Carlo simulations and for the model of protein adsorption.

## Pore formation

In this section we study the interaction between SNs and a free-standing bilayer and demonstrate that SNs with spikes long enough to pierce the lipid bilayer can induce reversible pores.

An horizontal free-standing bilayer was formed in a microfluidic chip at a desired location (see Materials and Methods). SNs of each type A, B, C, D were dispersed on one side in a buffer phase. SNs can be visualized and tracked by dark-field microscopy thanks to their plasmonic and scattering properties: the SNs are diffusing and bouncing on the bilayer surface, Fig. 2. The observation of SNs at the surface of the bilayer shows that none of these SNs are able to translocate through the lipid bilayer whatever the geometry of SNs. The average contact time of a SN with the bilayer is around a few seconds before SN detaches from the bilayer and moves in the solution. The SNs do not aggregate on the bilayer surface and are homogeneously distributed at the surface except at the edges of the bilayer, where we observe accumulation of SNs with non-negligible number of immobile SNs. These are clusters of SNs, which becomes much more visible for large concentrations of SNs, forming a golden ring at the edges of the bilayer, which is clearly seen under the microscope. The clustering at the edges demonstrates that the SNs are slightly hydrophobic, even if they are fully water-soluble and insoluble in oil. Such an effect can be related either to the presence of polyvinylpyrolidone on the surface of SN or to the Van der Walls interaction between the hydrophilic zwiterionic ligands coated on SNs (see Materials and Methods).

**Figure 2:**
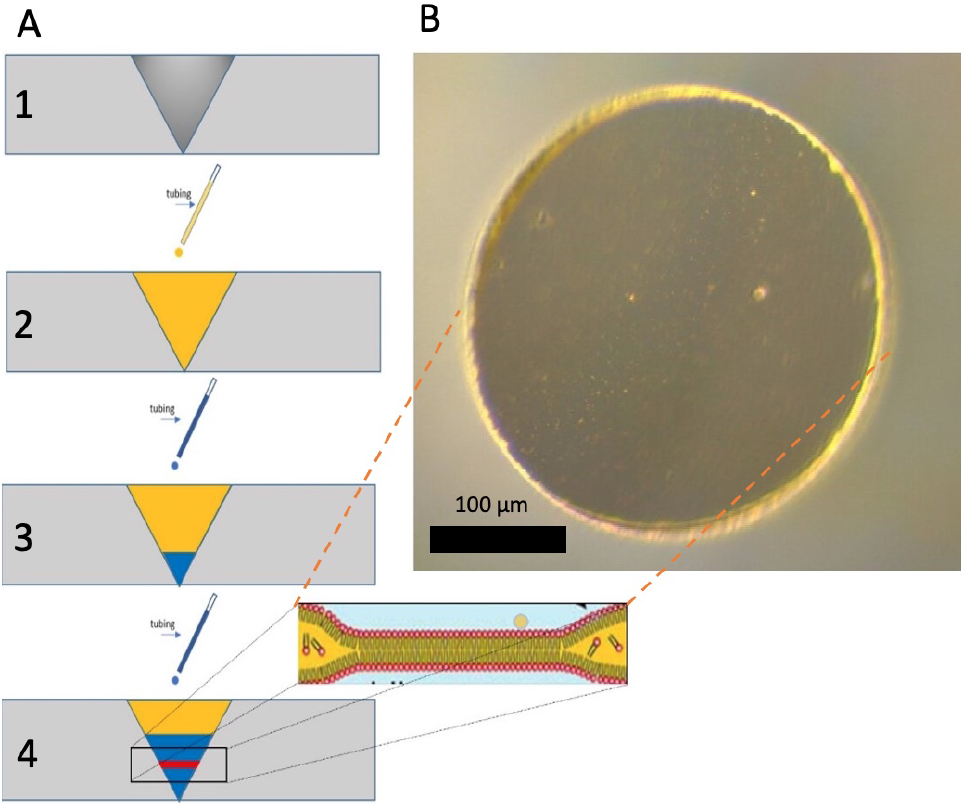
A) Schematic illustration of the bilayer formation from a mixture of lipid+oil (DOPC + Squalene) sandwiched between two water droplets: the bilayer forms after drainage of the oil, in 10-15 minutes.^35,36^ The resulting bilayer is stable for at least ≈ 0.5 - 2 hours. B) Visualization under dark-field microscopy of an horizontal lipid bilayer in the presence of spiky nanoparticles (100 nm diameter). SNs are depicted in orange, due to their plasmonic properties.

Microfluidic setup allows not only to directly visualize the interaction of nanoparticles with lipid bilayers, it permits simultaneously measure electrical signals across the membrane that are associated with interactions of individual nanoparticles.^37^ A planar lipid bilayer without inclusions and in the absence of nanoparticles represents a planar dielectric sheet, where its hydrophobic core is not permeable for ions from the buffer. Thus, if the voltage is applied between two sides of the bilayer, it represents a planar capacitor with nanometer-size thickness. Using a patch-clamp setup, one can measure a specific capacitance *C_s_* of this capacitor, which yields *C_s_* ≈ 3.9 mF/m^2^ for a DOPC free standing bilayer. Once the nanoparticles put in contact with the surface of the bilayer, the mechanical interaction changes the capacitance of the bilayer, thus providing a measurable electric signature of this interaction.

It is noteworthy to mention the influence on the measurements of the electric capacity of a golden ring formed at large concentrations of SNs (100 *μ*g gold/mL). For that purpose, SNs were dispersed via the bottom channel of the microfluidic device. Once the golden ring was formed at the edge of the bilayer, the bilayer was rinsed by diluting the surrounding phase around the bilayer by adding a buffer solution until the bulk concentration of SNs drops to zero, while SNs are present only at the bilayer edge. It turns out that the difference of the measured specific capacity *C_s_* with that of a pure DOPC lipid bilayer is negligible. Thus, one can conclude that the ring practically does not contribute to the measurements of the capacitance and the presence of the ring can be neglected. Nevertheless, the following part of our study corresponds to low concentrations of SNs where the ring does not form.

Following the dispersion of SNs in the bottom channel at ≈ 30 min after the bilayer formation, the capacitance and the conductance of the lipid bilayer in the presence of the SNs was measured as a function of time for different types of SNs, Fig. 3A-D). The specific capacitance of the bilayer is drastically reduced in presence of SNs, while the conductance of the bilayer exhibits massive increase, which is attributed to the poration of the bilayer by individual SNs. It is noteworthy that no significant conductance traces were measured without SNs. The size of the resulting nanopores formed by individual SN in contact with the bilayer can be determined by performing highly sensitive conductance measurements.

**Figure 3:**
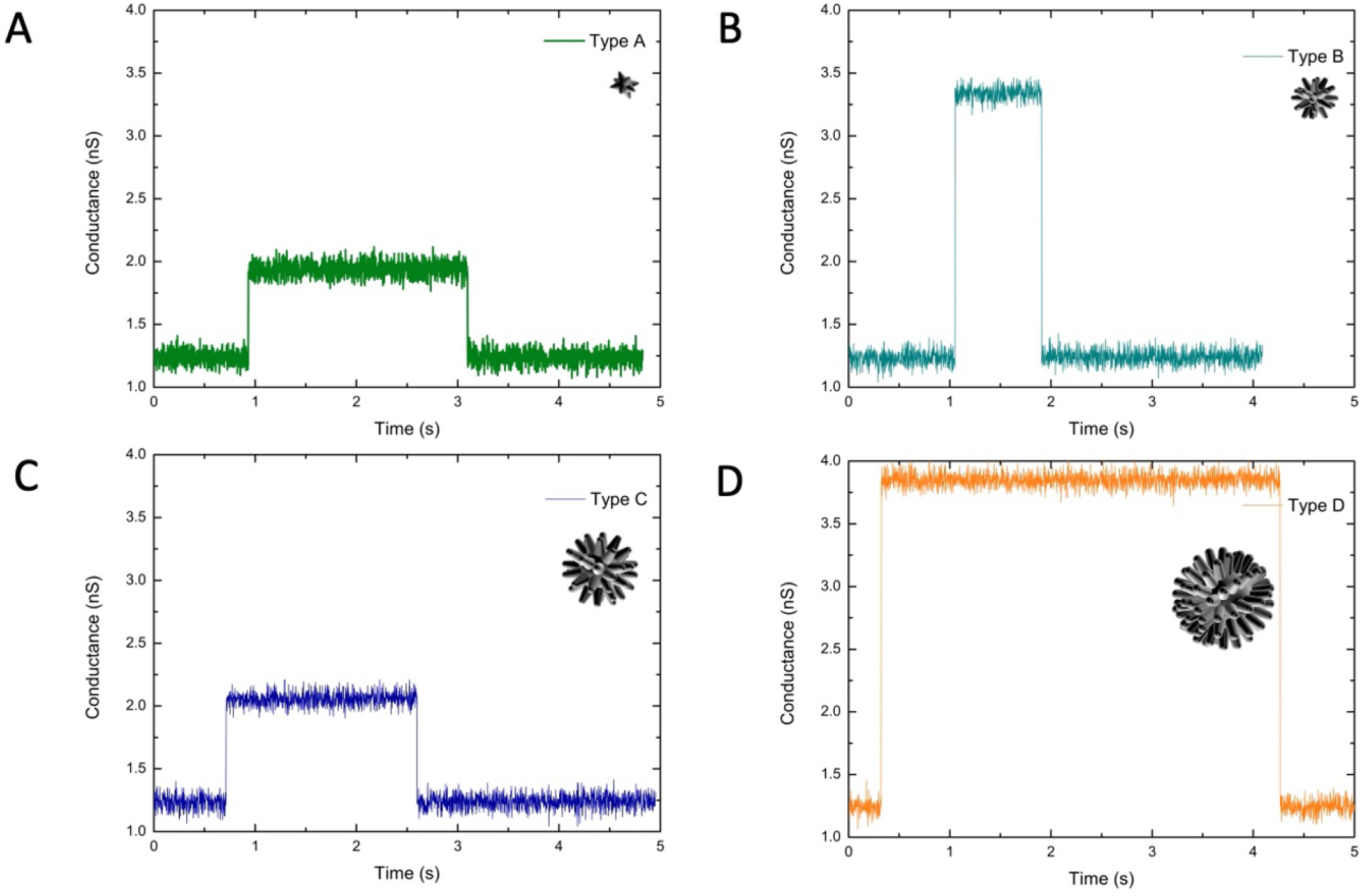
A)-D) Conductance traces from the poration of a DOPC bilayer in contact with SNs of Types A, B, C, D, correspondingly.

To this end, the conductance of the bilayer in the presence of SNs was measured at ultralow concentrations, 1 ng/ml. The corresponding conductance trace is showing well-defined conductance jumps with a characteristic conductance amplitude. A characteristic time of the jumps corresponds to the pore opening and closing events and indicates that the poration induced by SNs are reversible, while the amplitude is related to the the pore size. Indeed, the radius of the pore (in nm) is 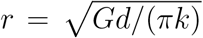, where *d* ~ 4 nm is the DOPC bilayer thickness, *k* = 1, 15 S/m is the bulk electrolyte conductivity (measured for 100 mM NaCl at 30°C), and *G* is the conductance (in nS).

We find that the size of the resulting pore *r* and its lifetime depend on the particular geometry of the SNs. For example, the biggest SNs of Type D induce long-lived (from 100 ms to a few seconds) conductance jumps with an amplitude corresponding to a pore radius of around ≈ 1.7 nm. Smaller SNs induce shorter-lived and smaller pores. The average radius of the pores is 0.88 for Type A, 1.52 for Type B, 0.95 for Type C, 1.7 nm for Type D. The size of the pores is not in the order of the size of SNs, which is gradually increasing from A to D, see Table 1, but most probably corresponds to the geometry of the spikes at the surface of the SNs. The 3D reconstruction of an average SN of each type suggests that Types A may contact with the bilayer with 3 or 4 spikes, while Types B and C may contact with 4 spikes and Type D with more than 4 spikes. This would suggest a gradual increase of pore sizes, however, the observed discrepancy in the order of the sizes could be attributed to the polydispersity of spike shapes that are formed randomly at the surfaces of the gold cores and spike sharpness. Nevertheless, the differences in the sizes is less than 1 nm, which is close to detection limit.

Noteworthy, the measured pores sizes correspond to the minimum pores sizes induced by SNs. Bigger pores lead to bilayer rupture and thus are not observed. Thus, these pore sizes can be considered pre-critical, *i.e.* before the bilayer rupture: the measured pore radii are close to the critical pore radius of rupture of the bilayer *r_c_* = *σ/*Γ ≈ 2.5 nm.^37^ *σ* ≈ 20 pN is the line tension of the pore edge corresponding to Γ ≈ 8 mN/m (see Materials and Methods). The pore line tension for a DOPC bilayer is estimated from Akimov et al. ^38^

Recently, the interaction of a spherical gold NPs coated with ZwBuEta with a lipid bilayer was reported.^36^ Based on electrophysiological experiments, it appears that these NPs do not create spontaneous pores for NPs diameters between 20 nm to 100 nm. This result can be used as a control for no spikes nanoparticles.

The difference between pores and the ability of an individual SN to break the bilayer is related to nanospikes geometry and hydrophobicity of the spikes. If the spikes are long enough and attractive, e.g. uniformly hydrophobic or hydrophilic, the bilayer would adsorb to reach the core of the spike and thus, then it has bigger chances to induce a rupture of the bilayer by area consumption. In contrast, if the spike is hydrophobic only at the tip or the spikes are not long, the SNs only anchoring the bilayer, making small pores without rupturing it. This was confirmed by Monte Carlo simulations using Bond Fluctuation Model ^39,40^ of a lipid bilayer interacting with three rigid spikes. The model of a lipid bilayer and the corresponding parameters are similar to Refs 41–43, where hydrophilic hydrophobic effects are mediated by short-range interactions. A hydrophobicity scale, *H_X_*, for a component “X” is defined relative to the lipid tail- and head group repulsion according Ref. 43 (Eq. (5) there). For a value of *H_X_* = 0, the component is hydrophilic, whereas for *H* = 1 the component is hydrophobic. The spikes are modeled as rigid and attractive truncated cones with spherical caps at the tips and variable distances between the centers, Fig. 4A). We define a total spike height of *h* = 52*a* with *a* being the lattice constant, a radius of the cap *R_c_* = 15*a*, and an opening angle *α* = 25°. The height of the corresponding full cone, *h*_0_, and the boundary height *h_b_* between conic and spherical part are given by

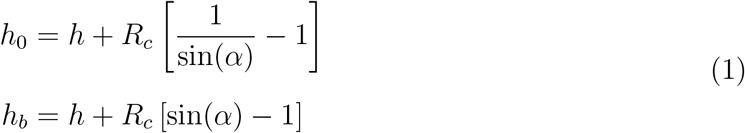

**Figure 4:**
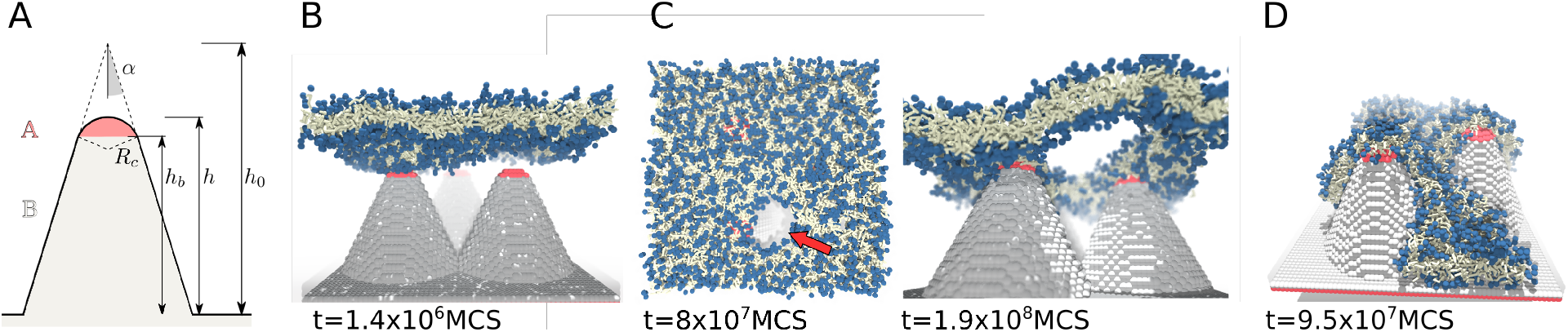
A) Spike geometry and notations in the coarse grained simulation model. Monte Carlo simulation snapshot of a bilayer for hydrophilicity *H_B_* = −0.16 (B,C) and *H_B_* = −0.32 (D) of the B-type substrate: B) initial contact with spikes, side view; C) pore formation (red arrow), top- and side views, typical pore radius for a period 8 × 10^7^ … 2 × 10^8^MCS was *r* = (22 ± 4)*a*; D) rupture of the bilayer due to full adsorption on the spikes for the stronger head-substrate attraction. The simulation time is given below each snapshot in units of Monte Carlo Steps (MCS).

Above the cap boundary height, *h_b_*, monomers are defines as hydrophobic (*H_A_* = 1). Below *h_b_*, monomers are defined super-hydrophilic *H_B_* = −0.16, i.e. only weakly attractive for the lipid head groups according to Eq. (5) in Ref. 43. Spikes are generated in the simulation box of cubic size 128*a* by occupying the volume with immobile monomers. Three spikes are positioned in an equilateral distance of 60*a* between the tips. The spikes are grafted to a substrate layer (immobile) with the same hydrophilic property *H_A_*. A number of 1200 lipids has been arranged in the simulation box in form of a bilayer above the spikes as well as an explicit solvent such that the volume occupancy in the mobile zones of the simulation box is 0.5. A fluctuation mode analysis for patches of 300 lipids with the same area per lipid revealed that the membrane is under slight tension of *γ* = 0.0142 *k_B_T/*(*a*^2^)^44^. Using an estimate, *a* ~ 0.154 nm^42^for the lattice constant, the tension can be referred to an order of magnitude near *γ* ~ 2.5 mN/m.

Since the diameter of the spikes tips is much larger than the bilayer thickness, the spikes do not pierce the bilayer, but due to adsorption of the bilayer on the spikes, the bilayer eventually becomes stretched in the zone between the spikes depending on the size of membrane-spike contact area. In addition, since pillar tips are hydrophobic, they are typically covered by lipid monolayer patches, that are connected to the remaining bilayer fraction between the tips. A defect in form of a boundary between monolayer and bilayer may act as a nucleation point for pores. Depending on the interplay between stretching, induced defects and bilayer fluctuations, a pore may form in the zone between spikes, Fig. 4B). In that respect, this mechanism is similar to the mechanism of pore formation described for nanostructured surfaces.^45^ Similarly, strong adsorption leads to the rupture of the lipid bilayer, see Fig. 4(D), where a larger surface-head attraction was applied by using *H_B_* = −0.32.

## Screening of spikes by protein corona

Once nanoparticles or biomaterials are introduced into biological media, within seconds they get coated by proteins.^46^ Dynamic exchange of proteins between the media and the surface of nanoparticles leads to the formation of PC, which is also changing with time.^47^ This process is very complex due to large diversity of proteins present in media, variability in kinetics and physico-chemical characteristics. PC not only interacts with the surfaces, it can also drastically modify the surface properties, screen the biomolecules present at the surfaces and screen or alter the action of ligands and coatings. To illustrate significant changes provoked by the adsorption of proteins in a well controlled microfluidic system, SNs of four types were placed in media with a model protein, bovine serum albumin, BSA. The coating with the protein lead to quantitative change of the interactions with lipid bilayers shown in previous section. Later, to confirm our findings that the protein effect is found not only for BSA protein, SNs were placed in blood serum media, containing large variety of proteins.

Addition of water-soluble BSA proteins to the solution of SNs leads to the formation of a protein corona. It is manifested in effective reduction of the length of spikes exposed to the interaction with lipid bilayers. To estimate the effect of BSA on a complex nanostructured surface of SNs of each type and calculate the maximal load of proteins per spike and per SN, we used the same approach that was applied for protein adsorption on planar nanostructured surfaces with spikes and pillars. ^46^ The kinetics of protein adsorption was modeled assuming irreversible adsorption of BSA protein, represented as a sphere of 7 nm diameter, on a shape of spike geometry given in Table 1 for each type of SN. Within Random Sequential Adsorption (RSA) approximation,^48^ the adsorption kinetics is expressed as a fraction of covered area *θ* (occupancy) as a function of time *t* via

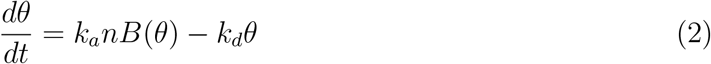

In this equation the blocking function *B*(*θ*) is defined as the probability to adsorb a protein to the surface for a given occupancy *θ*, while *k_a_* and *k_d_* are the rates of adsorption and desorption, respectively and *n* is the concentration of proteins in the bulk. The pristine surface *θ* = 0 corresponds to the blocking function *B*(0) = 1. Assuming irreversible adsorption of proteins within RSA approximation, the blocking function for a given geometry can be accessed directly with Monte Carlo simulations as *B*(*θ*) = *N_succ_/N_tot_*, where *N_succ_* is the number of potentially successful attempts and *N_tot_* is the total number of attempts.^46^ Using Gaussian approximation for the spikes shape

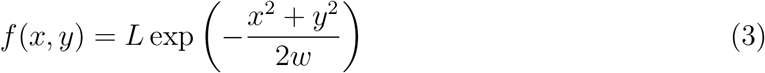

where *L* is the length of a spike, *w* is the Gaussian variance, related with the width of the spike. The results are present in Fig. 5A) for spikes and 5B) for SNs using the geometrical parameters of Table 1. It gives an estimate of a maximal number of proteins per spike and per SN. The protein load follow the order A-B-C-D, which is also consistent with their sizes and number of spikes per SN. This load leads to effective reduction of the spikes lengths and makes the SNs more round-shaped.

**Figure 5:**
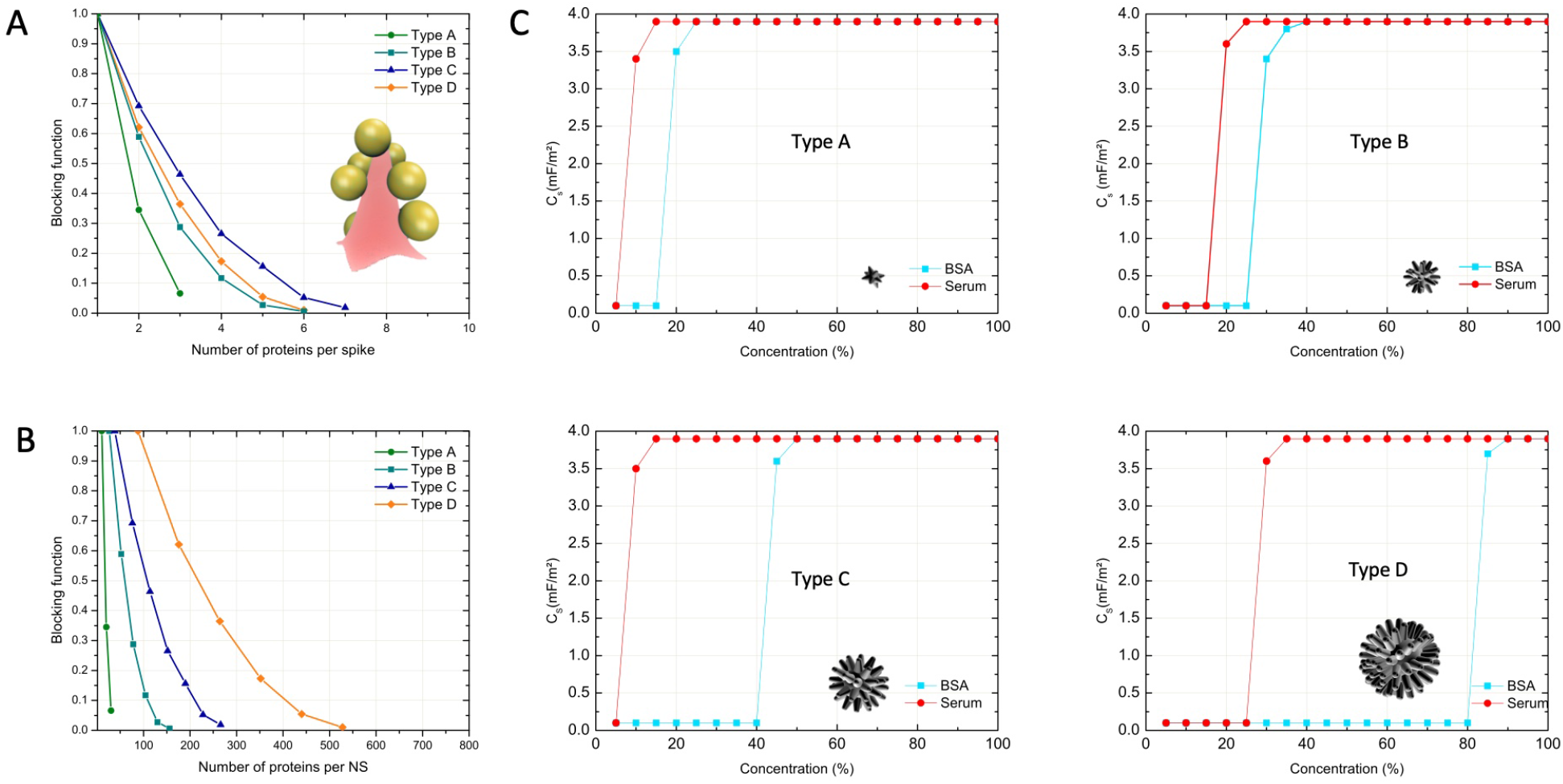
A) Bloking function of BSA proteins adsorbed on a spike of model SNs of types A-D, with the spike geometries described in Table 1. Inset: Illustration of a maximum load of BSA on Type A spike. B) Blocking function of BSA proteins per SN of Types A-D. They are deduced from A) and the number of spikes per partice. C) Specific capacitance of a DOPC bilayer in contact with the solution of SNs with BSA (blue) or with serum (red) as a function of BSA or serum concentration.

Microfluidic experiment confirms that addition of sufficiently high concentration of BSA proteins to the solution with SNs suppresses the pore formation. The formation of PC on the surface of SNs hinders the spikes and this results in the recovery of the bilayer original insulator properties similar to a pure lipid bilayer, regardless the presence of SNs on its surface. As expected, the screening effect of PC depends on both the geometry of the SNs and the BSA concentration. At a high protein concentration, above a threshold value for each type of SN, the pore formation is suppressed for all types of SNs. Nevertheless, the geometry of the spikes strongly influences the threshold value of the BSA concentration when the pores are no longer observed, Fig. 5C).

Another observation is that the pore suppression happens in a narrow concentration range close to the threshold value. This is probably due to the fact that the pores are created when the SNs are rolling on the surface and only by spikes in direct contact with the bilayer. Thus, although the accumulation of proteins can be a gradual process as a function of the bulk protein concentration, the suppression is happening only when all spikes are covered on all sides, which corresponds to the threshold value. The values of threshold concentrations follow the order A-B-C-D, which is consistent with the evaluation of maximal protein load from RSA. SNs of Type A do not induce pores in the bilayer at BSA bulk concentrations slightly below 20% and the threshold concentration for SNs of Type B is 25%. This is supported by electrophysiological measurements as no conductance traces are measured. SNs of of Type C and Type D require higher concentrations of BSA in solutions, 40% for Type C and 85% for Type D since they have larger areas and larger number of spikes to cover with proteins and these values are very high.

Thus, RSA estimations together with this result indicates that formation of PC around such nanoparticles is mainly determined by the geometry of nanoparticles, namely by the surface area and the number of spikes.

However, the threshold concentrations are much lower for serum proteins, red curves in Fig. 5C). This is due to the fact that serum contains large number of proteins of different types, that can fill the gaps between the spikes and at the surface of SNs more efficiently than globular-shaped BSA. Slightly hydrophobic surfaces of water-soluble SNs may also diminish serum protein adsorption. SNs of Type A and Type C, suppress pores formation when the buffer is containing 10% of serum. SNs of Types B and Type D, suppress pores at 20% and 30% of serum, correspondingly. It results that the spike geometry may hinder proteins adsorption and, thus, the formation of protein corona.

The geometry of the spikes strongly influences the threshold values of serum concentration leading to pores suppression. In particular, the sharpest spikes, *i.e.* SNs of type B and D, require higher serum concentration to recover the standard bilayer dielectric properties and thus, the interaction between the SNs and the bilayer is highly dependant on serum concentration. If we use the analogy of SNs with virus particles, that have the same sizes, one may expect that the serum and other proteins can screen the viruses from the interaction with bilayers, which may lead to various effects, *e.g.* escape from recognition by immune cells. It would be interesting to explore such effects with viral particles in separate experiments.

## Materials and Methods

### Synthesis of nanoparticles

4 types of gold spiky nanoparticles (SNs) with different core size and spike lengths (Types A, B,C, D, Fig. 1) were synthesized following a modification of the protocol described elsewhere.^32^ SNs of types A and B were made with 3 nm gold seeds, which were first formed by adding 22 *μ*L of HAuCl_4_ · 3 H_20_ (100 mM) to a PVP (MW 10.000; 36 *μ*M; 47.5 mL) solution prepared in the mixture DMF/water (v/v= 18/1). After 5 min under vigorous stirring, a freshly prepared NaBH_4_ aqueous solution (10 mM; 2.5 mL) was added quickly to the previous solution, which changed quickly from yellowish to pale pink color and kept stirred for another 2 hours. Solution was stored 24 hours before use. Type A and B differed only by the amount of 3 nm gold seeds added to the growth solution. In a typical synthesis, 82 *μ*L of HAuCl_4_ · 3 H_20_ (50 mM) was added to 15 mL of PVP solution (Mw 10.000; 10 mM) in DMF. Following this procedure, either 20 *μ*L or 5 *μ*L of the 3 nm gold seeds sol was added to the growth solution to prepare Au SNs type A and type B respectively. SNs were obtained within 2 hours with color change from yellowish to dark blue. SNs were washed 3 times to remove the PVP in excess and resuspended in water at 250 *μ*g Au/mL.

SNs Types C and D were made with 15 nm gold seeds, which were prepared by the citrate method adding 5 mL of citrate solution (1 wt%) to a boiling aqueous solution of HAuCl_4_ (100 mL, 0.5 mM) and allowed to react for 15 min and then cooled down at room temperature. Following this procedure, 5 mL of PVP (Mw 10.000; 0.1 mM) was added drop-wise to the previous solution and allowed to react overnight. The particles were then centrifuged at 8000 rpm for 30 min and redispersed in ethanol. Using the same protocol to prepare SNs type A and B, 82 *μ*L of HAuCl_4_ · 3 H_20_ (50 mM) was added to 15 mL of PVP solution (Mw 10.000; 10mM) in DMF. Following this, either 50 *μ*L or 15 *μ*L of the 15 nm gold seeds was added to the growth solution to prepare SNs Type C and Type D, respectively. SNs were obtained within 2 hours with color change from yellowish to blue/grey. SNs were washed 3 times to remove the PVP in excess and resuspended in water at 250 *μ*g Au/mL.

### Nanoparticle coating

All type of gold nanoparticles were coated with a thioctic bidentate sulfobetaine zwitterionic molecule ^49^ (Zw, 412 *μ*g/mol) to provide the same surface chemistry and known to enhance colloidal stability. ^50,51^ Briefly, in 10 mL of gold particles (100 *μ*g Au/mL), 40 *μ*L of NaOH (1 M) was added following by 100 *μ*L of Zw (0.5 mM) and stirred for 6 hours. Zw-coated particles were washed three times using centrifugation method (10.000 rpm for 20 min) to remove the free Zw. Sols were stored in water at 250 *μ*g Au/mL.

### Transmission Electron Microscopy (TEM)

Au SNs images were obtained on an FEI Tecnai G2 Twin TEM at 200 kV. TEM samples were prepared by placing a drop of SNs solution onto a copper grid covered with holey carbon films. Absorption spectra over a 190-900 nm range were collected using a Cary 100Bio UV-visible spectrophotometer (Varian).

### Microfluidic setup for horizontal bilayer formation

The formation of DOPC free-standing lipid bilayer in squalene oil was done as following. A PDMS 3D-chip was produced with a thin rectangular channel connected with a conical hole, made with a 3D printed method as detailed in.^35,36,52^ The bottom micro-channel of the PDMS chip was filled with a drop of oil+lipid followed by the delay of 5 min for spreading. Then a water droplet was added to the conical hole. The oily phase into the bottom channel was replaced by a water phase following the delay for the bilayer formation. The time to form the bilayer depends on the volume of the oil that separate the two water phases. Drainage of the oil happens in 1-2 hours, after that period of time the bilayer forms. Bilayer ideally forms at the intersection point where the upper conical wall and the bottom channel meet. The resulting formed bilayer is stable at least for 2 hours. The *μ*–chip was then placed under a dark-field microscope (Leica DM2700) for visualization with the camera DFC450 (Leica).

### Surface tension and bilayer tension measurements

Surface tensions of the various lipid monolayers at the oil/water interfaces were measured with the pendant drop method using a commercial measurement device (OCA 20, DataPhysics Instruments GmbH, Filderstadt, Germany). An oil solution with a concentration of 5 mg/ml DOPC was produced by introducing a droplet from a steel needle into the surrounding oil phase. The shapes of all droplets were fitted with the Young–Laplace equation to obtain their interfacial tension. After the initial formation of a droplet, the DOPC lipids adsorb to the interface, leading to a reduction in interfacial tension. This decrease was recorded over several minutes until a plateau was reached. From the values of the surface tension and the bilayer contact angle *θ*, which were obtained from pendant drop measurements or optical micrographs, respectively, the bilayer tension can be calculated using the Young equation:

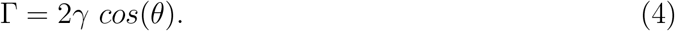

For a DOPC bilayer formed at a squalene oil/water interface, the associated bilayer tension could be estimated between Γ ≈ 8 mN/m. Nevertheless, it is worth to mention that this value may vary from a DOPC batch to another.

### Patch Clamping

Ag/AgCl electrodes were prepared by inserting a platinium electrode (HEKA, germany) in a borosilicate glass pipette (outer diameter 1.5 mm, inner diameter 0.86 mm, Vendor) containing an electrolyte agarose solution. Lipid membrane conductance was measured using the standard function provided by the patch clamp amplifier EPC 10 USB (Heka-Electronics). A 10 mV sinusoidal wave with a frequency of 20 kHz was used as an excitation signal. The electrodes are carefully introduced into the aqueous compartment of the Sylgard 184 device using micromanipulator. The specific capacitance *C_s_* = *C/A* is define by the ratio between the total capacitance *C* and the bilayer area *A*.

## Conclusions

On the example of spiky nanoparticles interacting with lipid bilayer we demonstrate how protein corona can change qualitatively the behavior of nanoparticles. Four types of water-soluble gold SNs coated with zwitterionic molecules with different core size and spikes geometries were synthesized. The geometry of synthetized SNs has a high a large surface-area-to-volume ratio and high curvature such that the nano-spikes. As a result, SNs dispersed near a lipid bilayer induce nanopores, which dimensions appear to be directly linked to SNs geometry. Monte Carlo simulations show that the pores are formed between the spikes anchoring into the bilayer, while the adsorption of the lipids at the surface of the spikes leads to the stretching of the bilayer between the spikes open a pore.

The presence of proteins in the solution change the behavior of SNs. PC formed around SNs hinders the effect of nanoparticle shape by reducing effectively the length of spikes and when the protein concentration reaches the threshold value, SNs do not induce pores. The threshold concentration appears to be directly linked to spikes geometry, which have dimensions comparable with the size of the proteins.

## Authors Contribution

All the authors designed the research. J.-B.F performed the microfluidic experiments. X.L.G performed the particles synthesis and their characterization. M.W. performed the Monte-Carlo numerical simulations. V.A.B. developed random sequential adsorption theory and performed the 3D reconstitution. All the authors analyzed their data and discussed the results. All the authors wrote the manuscript.

## Acknowledgments

J.-B.F acknowledges funding from SFB 1027 (DFG), project B4.

## Graphical TOC Entry

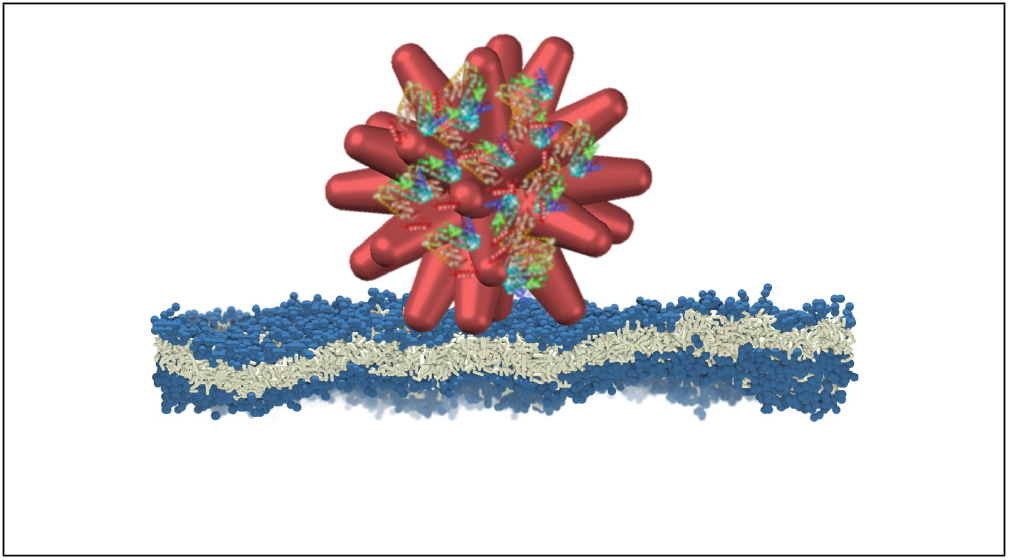

## References

(1) Patra, J. K.; Das, G.; Fraceto, L. F.; Campos, E. V. R.; Rodriguez-Torres, M. d. P.; Acosta-Torres, L. S.; Diaz-Torres, L. A.; Grillo, R.; Swamy, M. K.; Sharma, S.; Habtemariam, S.; Shin, H.-S. Nano based drug delivery systems: recent developments and future prospects. Journal of Nanobiotechnology 2018, 16, 71.

(2) Singh, R.; Lillard, J. W. Nanoparticle-based targeted drug delivery. Experimental and Molecular Pathology 2009, 86, 215–223.

(3) Nune, S. K.; Gunda, P.; Thallapally, P. K.; Lin, Y.-Y.; Forrest, M. L.; Berkland, C. J. Nanoparticles for biomedical imaging. Expert Opin Drug Deliv 2009, 6, 1175–1194.

(4) Brigger, I.; Dubernet, C.; Couvreur, P. Nanoparticles in cancer therapy and diagnosis. Advanced Drug Delivery Reviews 2012, 64, 24–36.

(5) Tran, S.; DeGiovanni, P.-J.; Piel, B.; Rai, P. Cancer nanomedicine: a review of recent success in drug delivery. Clin Trans Med 2017, 6, 44.

(6) Peer, D.; Karp, J. M.; Hong, S.; Farokhzad, O. C.; Margalit, R.; Langer, R. Nanocarriers as an emerging platform for cancer therapy. Nature Nanotechnology 2007, 2, 751–760, Number: 12 Publisher: Nature Publishing Group.

(7) De, M.; Ghosh, P. S.; Rotello, V. M. Applications of Nanoparticles in Biology. Advanced Materials 2008, 20, 4225–4241, eprint: https://onlinelibrary.wiley.com/doi/pdf/10.1002/adma.200703183.

(8) Werner, M.; Auth, T.; Beales, P. A.; Fleury, J. B.; Höök, F.; Kress, H.; Van Lehn, R. C.; Müller, M.; Petrov, E. P.; Sarkisov, L.; Sommer, J.-U.; Baulin, V. A. Nanomaterial interactions with biomembranes: Bridging the gap between soft matter models and biological context. Biointerphases 2018, 13, 028501, Publisher: American Vacuum Society.

(9) Lundqvist, M.; Stigler, J.; Elia, G.; Lynch, I.; Cedervall, T.; Dawson, K. A. Nanoparticle size and surface properties determine the protein corona with possible implications for biological impacts. Proc Natl Acad Sci U S A 2008, 105, 14265–14270.

(10) Oh, J. Y.; Kim, H. S.; Palanikumar, L.; Go, E. M.; Jana, B.; Park, S. A.; Kim, H. Y.; Kim, K.; Seo, J. K.; Kwak, S. K.; Kim, C.; Kang, S.; Ryu, J.-H. Cloaking nanoparticles with protein corona shield for targeted drug delivery. Nature Communications 2018, 9, 4548, Number: 1 Publisher: Nature Publishing Group.

(11) Mahmoudi, M.; Lynch, I.; Ejtehadi, M. R.; Monopoli, M. P.; Bombelli, F. B.; Laurent, S. Protein-Nanoparticle Interactions: Opportunities and Challenges. Chem. Rev. 2011, 111, 5610–5637, Publisher: American Chemical Society.

(12) Barbero, F.; Russo, L.; Vitali, M.; Piella, J.; Salvo, I.; Borrajo, M. L.; Busquets-Fité, M.; Grandori, R.; Bastús, N. G.; Casals, E.; Puntes, V. Formation of the Protein Corona: The Interface between Nanoparticles and the Immune System. Seminars in Immunology 2017, 34, 52–60.

(13) Cedervall, T.; Lynch, I.; Lindman, S.; Berggård, T.; Thulin, E.; Nilsson, H.; Dawson, K. A.; Linse, S. Understanding the nanoparticle–protein corona using methods to quantify exchange rates and affinities of proteins for nanoparticles. Proc Natl Acad Sci U S A 2007, 104, 2050–2055.

(14) Park, K. Facing the Truth about Nanotechnology in Drug Delivery. ACS Nano 2013, 7, 7442–7447, Publisher: American Chemical Society.

(15) Corbo, C.; Molinaro, R.; Parodi, A.; Toledano Furman, N. E.; Salvatore, F.; Tasciotti, E. The impact of nanoparticle protein corona on cytotoxicity, immunotoxicity and target drug delivery. Nanomedicine (Lond) 2016, 11, 81–100.

(16) Digiacomo, L.; Pozzi, D.; Palchetti, S.; Zingoni, A.; Caracciolo, G. Impact of the protein corona on nanomaterial immune response and targeting ability. WIREs Nanomedicine and Nanobiotechnology 2020, 12, e1615, eprint: https://onlinelibrary.wiley.com/doi/pdf/10.1002/wnan.1615.

(17) Ke, P. C.; Lin, S.; Parak, W. J.; Davis, T. P.; Caruso, F. A Decade of the Protein Corona. ACS Nano 2017, 11, 11773–11776, Publisher: American Chemical Society.

(18) Gulati, N. M.; Stewart, P. L.; Steinmetz, N. F. Bio inspired shielding strategies for nanoparticle drug delivery applications. Mol Pharm 2018, 15, 2900–2909.

(19) Pino, P. d.; Pelaz, B.; Zhang, Q.; Maffre, P.; Nienhaus, G. U.; Parak, W. J. Protein corona formation around nanoparticles – from the past to the future. Mater. Horiz. 2014, 1, 301–313, Publisher: The Royal Society of Chemistry.

(20) Pelaz, B.; del Pino, P.; Maffre, P.; Hartmann, R.; Gallego, M.; Rivera-Fernández, S.; de la Fuente, J. M.; Nienhaus, G. U.; Parak, W. J. Surface Functionalization of Nanoparticles with Polyethylene Glycol: Effects on Protein Adsorption and Cellular Uptake. ACS Nano 2015, 9, 6996–7008, Publisher: American Chemical Society.

(21) Schöttler, S.; Becker, G.; Winzen, S.; Steinbach, T.; Mohr, K.; Landfester, K.; Mailänder, V.; Wurm, F. R. Protein adsorption is required for stealth effect of poly(ethylene glycol)- and poly(phosphoester)-coated nanocarriers. Nat Nanotechnol 2016, 11, 372–377.

(22) Visalakshan, R. M.; García, L. E. G.; Benzigar, M. R.; Ghazaryan, A.; Simon, J.; Mierczynska-Vasilev, A.; Michl, T. D.; Vinu, A.; Mailänder, V.; Morsbach, S.; Landfester, K.; Vasilev, K. The Influence of Nanoparticle Shape on Protein Corona Formation. Small 2020, 16, 2000285, eprint: https://onlinelibrary.wiley.com/doi/pdf/10.1002/smll.202000285.

(23) García-Álvarez, R.; Hadjidemetriou, M.; Sánchez-Iglesias, A.; Liz-Marzán, L. M.; Kostarelos, K. In vivo formation of protein corona on gold nanoparticles. The effect of their size and shape. Nanoscale 2018, 10, 1256–1264, Publisher: The Royal Society of Chemistry.

(24) Vinluan, R. D.; Zheng, J. Serum protein adsorption and excretion pathways of metal nanoparticles. Nanomedicine (Lond) 2015, 10, 2781–2794.

(25) Nguyen, V. H.; Meghani, N. M.; Amin, H. H.; Tran, T. T. D.; Tran, P. H. L.; Park, C.; Lee, B.-J. Modulation of serum albumin protein corona for exploring cellular behaviors of fattigation-platform nanoparticles. Colloids Surf B Biointerfaces 2018, 170, 179–186.

(26) Sloan-Dennison, S.; Schultz, Z. D. Label-free plasmonic nanostar probes to illuminate in vitro membrane receptor recognition †Electronic supplementary information (ESI) available: Experimental details, Fig. S1–S9, and Table S1. See DOI: 10.1039/c8sc05035j. Chem Sci 2018, 10, 1807–1815.

(27) Chang, Y.-X. et al. Gold Nanotetrapods with Unique Topological Structure and Ultranarrow Plasmonic Band as Multifunctional Therapeutic Agents. J. Phys. Chem. Lett. 2019, 10, 4505–4510, Publisher: American Chemical Society.

(28) Bosch, B. J.; Zee, R. v. d.; Haan, C. A. M. d.; Rottier, P. J. M. The Coronavirus Spike Protein Is a Class I Virus Fusion Protein: Structural and Functional Characterization of the Fusion Core Complex. Journal of Virology 2003, 77, 8801–8811, Publisher: American Society for Microbiology Journals Section: STRUCTURE AND ASSEMBLY.

(29) Ezzat, K. et al. The viral protein corona directs viral pathogenesis and amyloid aggregation. Nat Commun 2019, 10, 2331.

(30) Gunawan, C.; Lim, M.; Marquis, C. P.; Amal, R. Nanoparticle–protein corona complexes govern the biological fates and functions of nanoparticles. J. Mater. Chem. B 2014, 2, 2060–2083, Publisher: The Royal Society of Chemistry.

(31) Hadjidemetriou, M.; Kostarelos, K. Evolution of the nanoparticle corona. Nature Nanotechnology 2017, 12, 288–290, Number: 4 Publisher: Nature Publishing Group.

(32) Barbosa, S.; Agrawal, A.; Rodríguez-Lorenzo, L.; Pastoriza-Santos, I.; Alvarez-Puebla, R. A.; Kornowski, A.; Weller, H.; Liz-Marzán, L. M. Tuning size and sensing properties in colloidal gold nanostars. Langmuir 2010, 26, 14943–14950.

(33) Kumar, P. S.; Pastoriza-Santos, I.; Rodríguez-González, B.; Abajo, F. J. G. d.; Liz-Marzán, L. M. High-yield synthesis and optical response of gold nanostars. Nanotechnology 2007, 19, 015606, Publisher: IOP Publishing.

(34) Kedia, A.; Senthil Kumar, P. Precursor-Driven Nucleation and Growth Kinetics of Gold Nanostars. J. Phys. Chem. C 2012, 116, 1679–1686, Publisher: American Chemical Society.

(35) Heo, P.; Ramakrishnan, S.; Coleman, J.; Rothman, J. E.; Fleury, J.-B.; Pincet, F. Highly Reproducible Physiological Asymmetric Membrane with Freely Diffusing Embedded Proteins in a 3D-Printed Microfluidic Setup. Small 2019, 15, 1900725, eprint: https://onlinelibrary.wiley.com/doi/pdf/10.1002/smll.201900725.

(36) Linklater, D. P.; Baulin, V. A.; Guével, X. L.; Fleury, J.-B.; Hanssen, E.; Nguyen, T. H. P.; Juodkazis, S.; Bryant, G.; Crawford, R. J.; Stoodley, P.; Ivanova, E. P. Antibacterial Action of Nanoparticles by Lethal Stretching of Bacterial Cell Membranes. Advanced Materials 2020, 32, 2005679, eprint: https://onlinelibrary.wiley.com/doi/pdf/10.1002/adma.202005679.

(37) Guo, Y.; Terazzi, E.; Seemann, R.; Fleury, J. B.; Baulin, V. A. Direct proof of spontaneous translocation of lipid-covered hydrophobic nanoparticles through a phospholipid bilayer. Science Advances 2016, 2, e1600261, Publisher: American Association for the Advancement of Science Section: Research Article.

(38) Akimov, S. A.; Volynsky, P. E.; Galimzyanov, T. R.; Kuzmin, P. I.; Pavlov, K. V.; Batishchev, O. V. Pore formation in lipid membrane II: Energy landscape under external stress. Scientific Reports 2017, 7, 12509, Number: 1 Publisher: Nature Publishing Group.

(39) Carmesin, I.; Kremer, K. The bond fluctuation method: a new effective algorithm for the dynamics of polymers in all spatial dimensions. Macromolecules 1988, 21, 2819–2823.

(40) Deutsch, H. P.; Binder, K. Interdiffusion and self-diffusion in polymer mixtures: A Monte Carlo study. The Journal of Chemical Physics 1991, 94, 2294–2304.

(41) Werner, M.; Sommer, J. U. Polymer-decorated tethered membranes under good- and poor-solvent conditions. Eur. Phys. J. E 2010, 31, 383–392.

(42) Pogodin, S.; Werner, M.; Sommer, J.-U.; Baulin, V. A. Nanoparticle-Induced Permeability of Lipid Membranes. ACS Nano 2012, 6, 10555–10561.

(43) Werner, M.; Sommer, J.-U. Translocation and Induced Permeability of Random Amphiphilic Copolymers Interacting with Lipid Bilayer Membranes. Biomacromolecules 2015, 16, 125–135.

(44) Werner, M. Interaction of Polymers with Self-Assembled Lipid Bilayer Membranes : Translocation and Pore Formation at Balanced Hydrophobicity /. Dissertation 2016, Technische Universität Dresden.

(45) Pogodin, S.; Hasan, J.; Baulin, V.; Webb, H.; Truong, V.; Phong Nguyen, T.; Boshkovikj, V.; Fluke, C.; Watson, G.; Watson, J.; Crawford, R.; Ivanova, E. Biophysical Model of Bacterial Cell Interactions with Nanopatterned Cicada Wing Surfaces. Biophysical Journal 2013, 104, 835–840.

(46) Manzi, B. M.; Werner, M.; Ivanova, E. P.; Crawford, R. J.; Baulin, V. A. Simulations of Protein Adsorption on Nanostructured Surfaces. Scientific Reports 2019, 9, 4694, Number: 1 Publisher: Nature Publishing Group.

(47) Vroman, L. The life of an artificial device in contact with blood: initial events and their effect on its final state. Bull N Y Acad Med 1988, 64, 352–357.

(48) Schaaf, P.; Talbot, J. Surface exclusion effects in adsorption processes. J. Chem. Phys. 1989, 91, 4401–4409.

(49) Park, J.; Nam, J.; Won, N.; Jin, H.; Jung, S.; Jung, S.; Cho, S.-H.; Kim, S. Compact and Stable Quantum Dots with Positive, Negative, or Zwitterionic Surface: Specific Cell Interactions and Non-Specific Adsorptions by the Surface Charges. Advanced Functional Materials 2011, 21, 1558–1566, eprint: https://onlinelibrary.wiley.com/doi/pdf/10.1002/adfm.201001924.

(50) Susumu, K.; Oh, E.; Delehanty, J. B.; Blanco-Canosa, J. B.; Johnson, B. J.; Jain, V.; Hervey, W. J.; Algar, W. R.; Boeneman, K.; Dawson, P. E.; Medintz, I. L. Multifunctional compact zwitterionic ligands for preparing robust biocompatible semiconductor quantum dots and gold nanoparticles. J Am Chem Soc 2011, 133, 9480–9496.

(51) Porret, E.; Sancey, L.; Martín-Serrano, A.; Montañez, M. I.; Seeman, R.; Yahia-Ammar, A.; Okuno, H.; Gomez, F.; Ariza, A.; Hildebrandt, N.; Fleury, J.-B.; Coll, J.-L.; Le Guével, X. Hydrophobicity of Gold Nanoclusters Influences Their Interactions with Biological Barriers. Chem. Mater. 2017, 29, 7497–7506, Publisher: American Chemical Society.

(52) Tawfik, H.; Puza, S.; Seemann, R.; Fleury, J.-B. Transport Properties of Gramicidin A Ion Channel in a Free-Standing Lipid Bilayer Filled With Oil Inclusions. Front Cell Dev Biol 2020, 8, 531229.

